# Evidence for infection of *Eucalyptus camaldulensis* with a *Leifsonia* bacterium in Australia and Syria using high throughput sequencing

**DOI:** 10.1101/2024.07.16.603836

**Authors:** J.W. Randles, D. Hanold, U. Baumann, A. Borneman

## Abstract

A biotic hypothesis for the eucalypt disease of unknown aetiolgy, Mundulla Yellows (MY), was tested by seeking non-host nucleic acids in affected *Eucalyptus camaldulensis* trees using high throughput sequencing (HTS). Ten contigs were selected from a pool of nucleic acids unique to diseased trees, and oligonucleotide probes were prepared for specific dot-blot hybridisation assay of these contigs. One of the contigs was found to be associated with a few MY-affected trees collected in South Australia and Western Australia, as well as with similarly affected *E. camaldulensis* trees growing in Syria. The probes for this contig represented opposite polarities of the contig, and both hybridised with the same positive samples even after alkali treatment confirming that the contig that they represented was dsDNA. This contig, comprising 766 nt, showed 95% similarity to whole genome shotgun sequence of *Leifsonia xyli*. This preliminary microbiome analysis of MY affected trees provides the first evidence that a *Leifsonia* species may infect eucalypts and that it should be further studied as a potential agent of MY.

## INTRODUCTION

Attempts to determine whether the Australian eucalypt dieback and leaf-yellowing disease of unknown aetiology, described as Mundulla Yellows (MY) (Keane et al., 2000; Hanold et al., 2002) is biotic or abiotic have given contradictory results. In mature field trees, symptom descriptors include chlorosis, mosaic and distortion of leaves, a progressive slow die-back of affected limbs and epicormic shoot production (Hanold et al., 2006). Similar symptoms have been observed in eucalypts growing in other countries (Hanold and Randles, 2003; Hanold et al., 2002; 2006; 2009). *Eucalyptus camaldulensis* seedlings experimentally grafted with bark patches from symptomatic eucalypts show similar leaf symptoms, loss of apical dominance and dieback, implying involvement of a transmissible biotic agent (Hanold et al., 2006). No specific pathogen has been shown to cause MY, although associations of the nematode *Merlinius* spp, virus-like particles, and an 84kDa and 116kDa protein with the disease (Hanold et al., 2006; Luck et al., 2006; Randles et al., 2010) support the view that one or more infectious agents may contribute to disease expression. Abiotic nutritional and soil factors such as those inducing lime chlorosis (Luck et al., 2006; Czerniakowski et al., 2006) have been implicated in symptom development by the alleviation of foliar and crown symptoms with iron and manganese implants (Schultz and Good, 2018), but longterm recovery from the full disease syndrome indicating a single abiotic cause has not been reported. It is important to determine whether there is a consistent association between a principal biotic or abiotic factor so that management strategies can be developed as for other diseases of eucalypts (Keane et al., 2000).

We report the use of high throughput sequencing (HTS) with selected MY affected and unaffected *E. camaldulensis* trees to look for non-host nucleic acids that may differentiate diseased from normal trees. A disease-associated contig representing the bacterial genus *Leifsonia* was identified in both Australian and Syrian samples, and the implications of this finding are discussed.

## MATERIALS AND METHODS

### Samples

The trees used for this study were *E. camaldulensis* (except where indicated) growing in South Australia (SA), Western Australia (WA) and Syria (Table 1). Field trees from SA came from Adelaide (including the University of Adelaide Waite campus at Urrbrae, and Burnside) and Mundulla. C40, a commercially available *E. camaldulensis* line, derived from sterile tissue culture (synonymous with A40; Hanold et al., 2006) and maintained in sterilised potting soil under glasshouse containment conditions, was used as the disease-free negative control. Trees with characteristic MY symptoms (Hanold et al., 2006) were sampled at Mundulla. Trees from WA either showed symptoms like those of MY at Mundulla, or dieback alone. Trees in Syria came from the ICARDA (International Center for Agricultural Research in the Dry Areas) campus in Aleppo and had a foliar chlorosis disease previously described as resembling MY (Hanold et al., 2009).

**Table 1.**
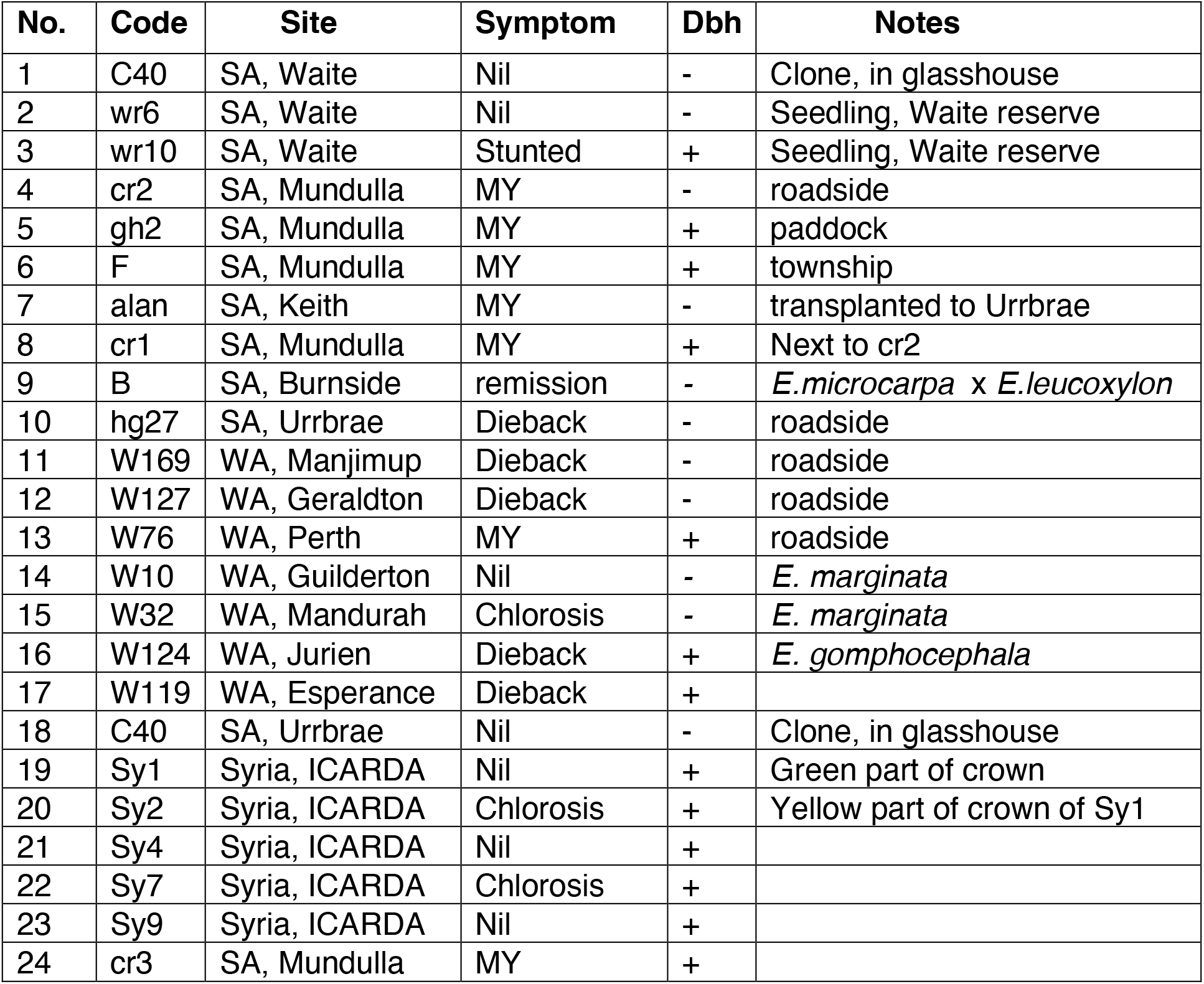
Trees identified by number, code, site, symptom descriptor, and dot-blot hybridisation (Dbh) results as shown in Figs 2 and 3. Except where indicated, all samples were from *E. camaldulensis*.

### Preparation of asymptomatic (H) and symptomatic (D) nucleic acid pools

Nucleic acids were extracted from young expanded leaves of three trees without MY symptoms (Table 1, trees 1-3; pool H) and three symptomatic trees (Table 1, trees 4-6; pool D). These pools (30g) comprised 10g from each of the three trees and were blended in 10 volumes of extraction buffer (0.4M NaCl, 0.5% monothioglycerol, 0.1M Tris-HCl, pH 7.2, 5mM EDTA, 10% phenol). Aqueous phenol (90% v/v) was added to 33% (v/v), sodium dodecyl sulphate (SDS) was added to 2%, and the resulting emulsion was stirred for 60 min at 22°C. Following centrifugation at 10,000g for 15 min, the supernatant was recovered and re-extracted with an equal volume of phenol/chloroform (1:1), then chloroform alone, by centrifugation. Nucleic acids were precipitated with an equal volume of isopropanol, dried, and the resulting pellet was resuspended and incubated in 2.5mL proteinase K (Promega) at 50ug/mL in 0.5%

SDS, 0.1M NaCl, 50mM Tris-HCl, pH 8, at 25°C for 14 hours. Following a single phenol extraction nucleic acids were precipitated with an equal volume of isopropanol, drained and dried. DNAse I (Promega RQ1; 2 units in 20 μL reaction buffer) was added to the pellet and incubated at 37°C for 60 min. Following a single phenol extraction, isopropanol precipitation in 0.1M sodium acetate, and a wash in ethanol, pellets were dried and resuspended in sterile water to give a final concentration of 2.4 μg/μL.

Ribosomal RNA was removed with a Ribominus kit (Invitrogen). The resulting extract was precipitated with ethanol (5 vol in 0.1M Na acetate), dried and adjusted to 0.48 μg/μL in water. Aliquots of 11 μL (5 μg) of each sample were transcribed to cDNA using random primers to generate unstranded libraries (Hrdlickova et al., 2016).

### High throughput sequencing (HTS)

Pools H and D were subjected to paired-end HTS using Illumina technology (AGRF, Australia). Sequence reads were examined with FastQC (Andrews, 2010) and subjected to adapter and quality trimming using the FASTX toolkit (Hannon, 2010). Over-represented sequences consisted of rRNA and were subsequently removed based on a BlastN sequence comparison to rRNA sequences of *Rosids* retrieved from the NCBI database (2011). The remaining reads for pool H (2.7 ×10^6^) and pool D (1.19×10^6^) were assembled by MIRA 3 (Chevreux et al., 2004). Of the 8648 contigs only those unique to the D pool were selected for further analysis. Contigs were used as queries in BlastN searches against a) *Eucalyptus* genomic sequences available at the time and b) NCBI’s nr-database (2011). Contigs virtually identical with eukaryotic sequences in either database were filtered out. The remaining contigs were considered as candidates.

### Preparation of probes

Ten of the most abundant contigs unique to pool D were selected, and pairs of representative oligonucleotide probes (26 to 51 nt in size) complementary to each polarity of the contig, and separated by at least 48 nts from each other, were synthesised (GeneWorks, Adelaide). The Tm of each pair was matched to within 5^0^ C. They were end-labelled using *γ*32P-ATP and polynucleotide kinase with the Promega DNA 5’ End-Labeling System. Probes were denatured in 50% formamide at 80°C then added to hybridisation buffer (Hanold and Randles, 1991).

### Preparation of field samples for dot-blot hybridisation

Frozen (1g) or dehydrated (0.3g) leaf samples were ground to a powder in liquid nitrogen with a pestle and mortar. Five mL of extraction buffer (100mM Tris-HCl, pH 7.3, 100mM sodium acetate, 400mM NaCl, 10mM EDTA, 2% SDS) containing 0.5% monothioglycerol and 0.2g of polyvinylpyrrolidone was added with mixing at room temperature for 45min, 0.5 vol of chloroform was added and mixing continued for 5min. Following centrifugation (ca. 7500g), the aqueous phase was recovered and extracted with an equal volume of 90% aqueous phenol for 45 min. After centrifugation nucleic acids were recovered from the upper phase by precipitation with an equal volume of isopropanol. Pellets were dried and dissolved in 50μL of water. Each sample was applied as 1μL dots in a 10-fold dilution series to nylon (Zetaprobe, BioRad) membrane. Replicates of panels representing the samples were fixed by UV irradiation for hybridisation assay.

### Hybridisation assay

Membranes were washed in 0.1xSSC, 0.1% SDS at 67°C for 1h, prehybridised in hybridisation buffer for 16h (Hanold and Randles, 1991) then incubated with one of the probe specific hybridisation mixtures at 42°C for 21h. They were rinsed with 0.5xSSC, 0.1% SDS at 20°C, then washed in 1xSSC, 0.1%SDS, 55°C for 1h before autoradiography.

To test for alkali resistance of the fixed samples, replicate membranes were pre-incubated in 0.3M KOH for 1 h at 22°C and washed in distilled water before hybridisation as above.

## RESULTS

### Comparison of ten oligonucleotide probes for differentiating samples

Of the ten pairs of oligonucleotide probes representing the ten most abundant pool D (diseased trees) specific contigs, only one pair, named here as 14A and 14B after the contig 14 they represented, hybridised significantly to any of the samples. The contig with their position is shown in Fig. 1.

**Figure 1.**
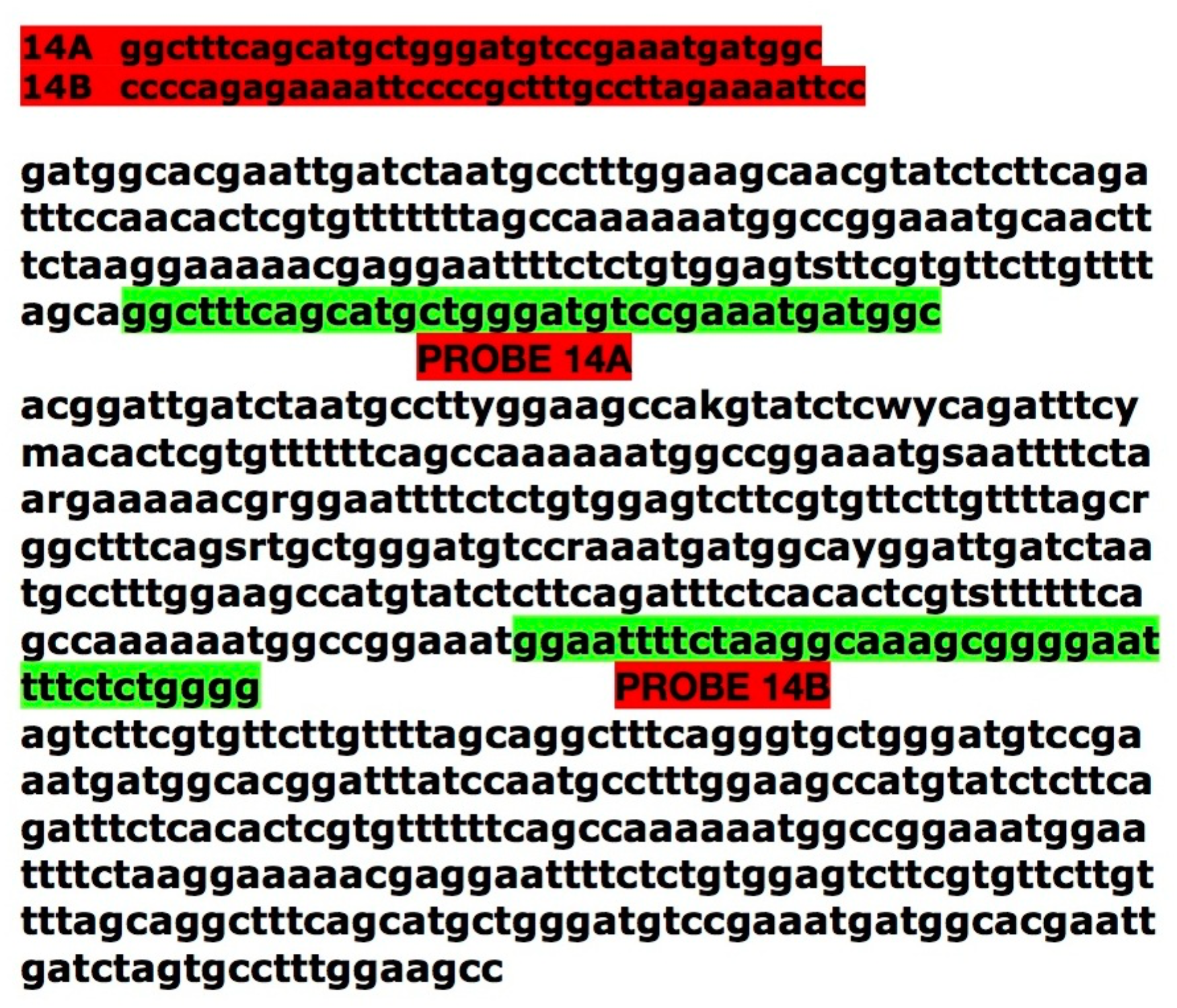
Sequence of the pool D-associated contig 14, shown in one of two alternative orientations. Oligonucleotide probe sequences 14A and 14B are shown in red; target complementary sequences are shown in green. Thus, probe 14B was used to hybridise to and identify the sequence shown, whereas probe 14A was used to detect the complementary strand.

### Association between symptom presentation and hybridisation to probes representing contig 14

Fig. 2 shows the semiquantitative dotblot hybridisation (Dbh) assay results. Table 1 summarises them according to site, species, and symptoms observed.

**Figure 2.**
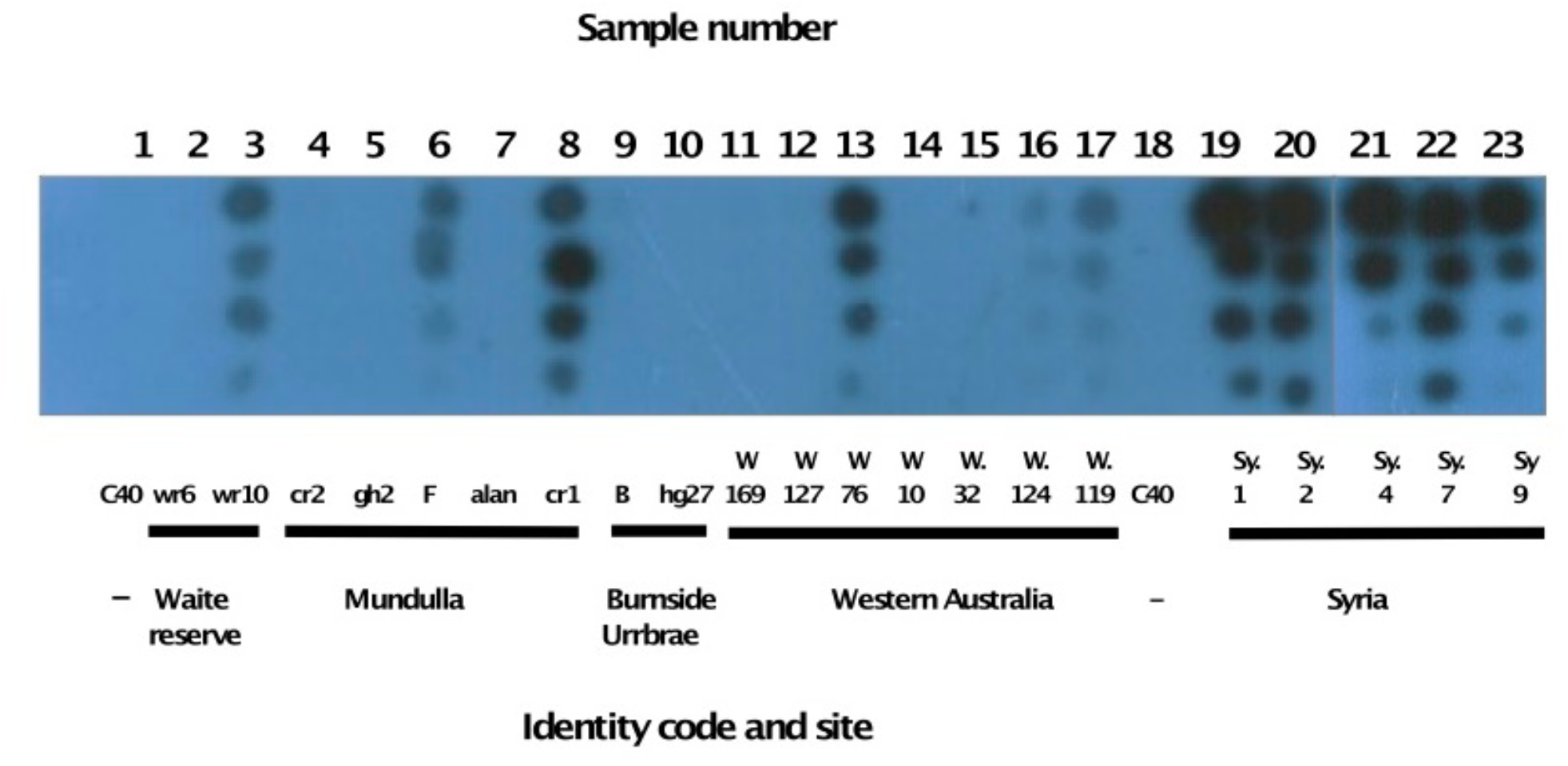
Autoradiograph of a semiquantitative dot-blot hybridisation assay with oligonucleotide probe 14A against the samples listed in Table 1. From the top, each 1μL sample was spotted undiluted and diluted 1:10, 1:100 and 1:1000. (Note that gh2 was negative here, but positive in Fig. 3).

The asymptomatic control samples from SA (trees 1, 2, 18) were negative. Four trees (5,6,8 and 24) with MY symptoms and one with stunting (3) were positive. Two trees with dieback or remission from early symptoms (9, 10) were negative.

In WA, one *E. camaldulensis* tree with MY symptoms (13) and one (17) with dieback were positive. Another species with dieback, *E. gomphocephala*, was positive, while two *E*.*marginata* trees were negative.

Of the four trees sampled in Syria, all were positive, whether they had chlorosis symptoms or not.

Thus, there was a strong association between positive hybridisation and the expression of MY symptoms. In addition, some trees with dieback, chlorosis or stunting were also positive.

### Evidence that probes had bound to a double-stranded DNA target

To test whether the probes had hybridised to an RNA or DNA target sequence, we applied alkali hydrolysis to a pair of membranes before hybridisation. Probes 14A and 14B both hybridised to the same samples (pool H, cr1, W76, gh2, cr3, and Sy2) before and after alkali treatment (Fig. 3) while the healthy control (C40) was negative. The positive signal with pool H was due to the inclusion of the stunted but positive tree wr10 in the three samples comprising this pool. Pool D was also positive (not shown). These results together show that the positive samples were double stranded DNA.

**Figure 3.**
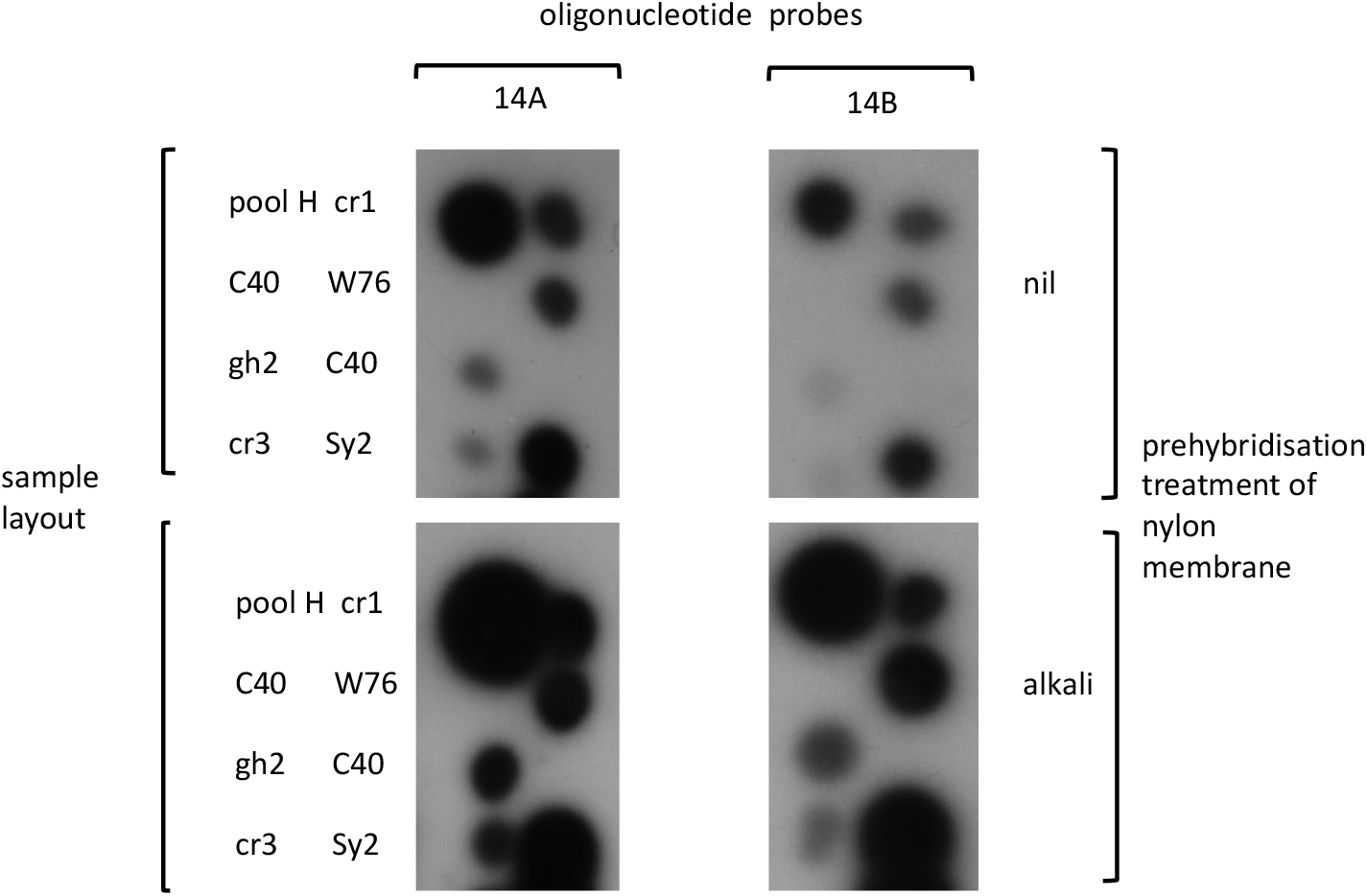
Autoradiographs of four nylon membrane replicates (sample layout shown on the left). The upper pair of membranes was untreated(nil), the lower pair was pretreated with alkali prior to hybridisation (description on right). The two membranes on the left were hybridised with oligonucleotide probe 14A, the two on the right with 14B.

### Homology of contig 14 with *Leifsonia xyli*

Fig. 4 shows that the 766 nt contig 14 has ca. 95% identity at the DNA level with *Leifsonia xyli* subsp. *xyli* gdw1 scaffold 43, whole genome shotgun sequence in GenBank (NCBI GenBank Acc. No. LNZG01000038.1). The oligonucleotide probes used to detect this contig showed 89% (14A) and 92% (14B) similarity to their respective targets.

**Figure 4.**
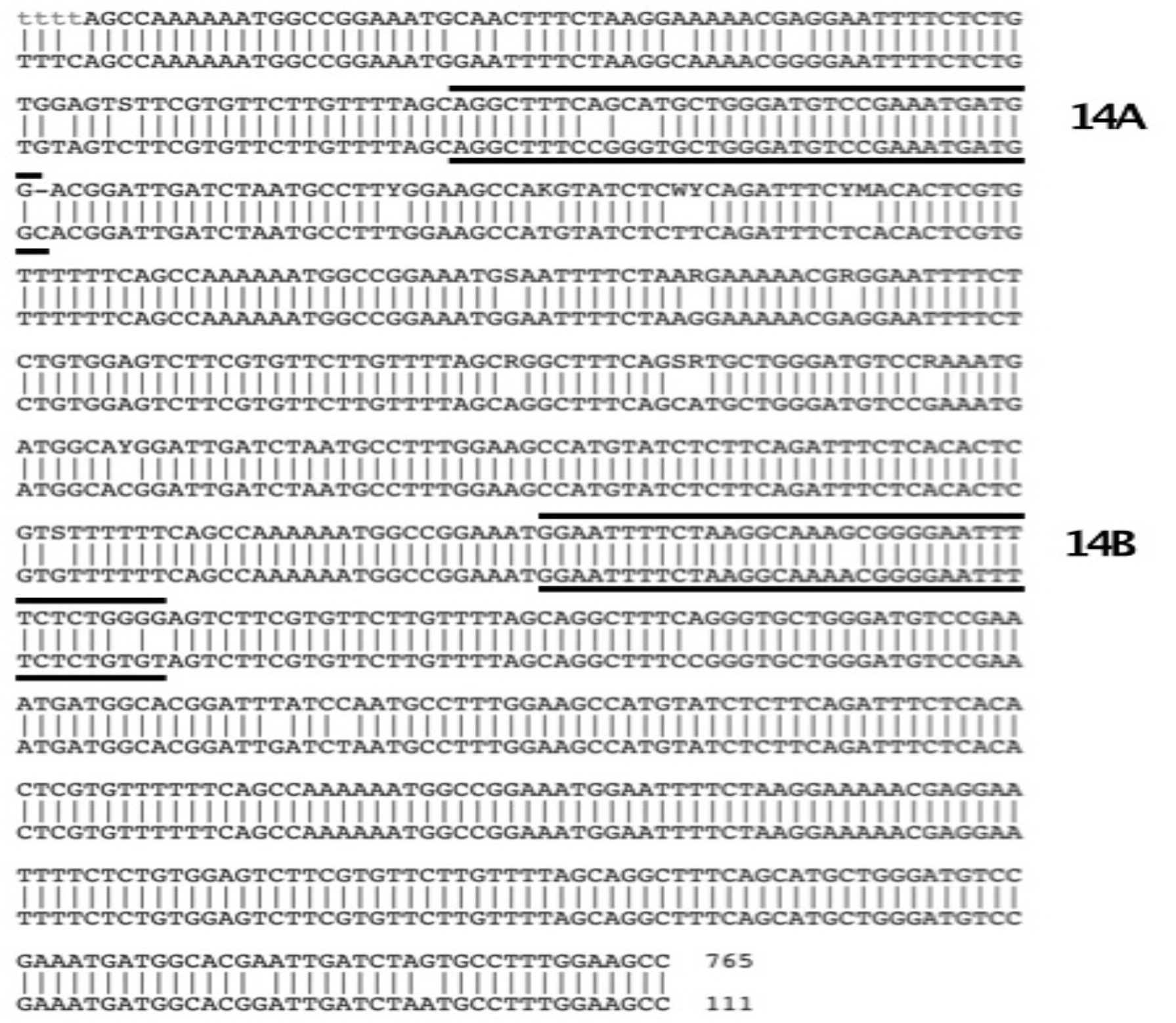
GenBank nucleotide Blastn comparison of contig 14 (upper) with whole genome shotgun sequence LNZG01000038.1 in the NCBI GenBank database. The targets of oligonucleotide probes 14A and 14B are indicated with bold lines.

## DISCUSSION

We describe the use of Illumina HTS to seek a putative biotic agent for MY by identifying genomic components foreign to the eucalypt host. A comparison of contigs constructed from our pools of asymptomatic and symptomatic trees identified a few which were potentially associated with symptoms of the disease. None of these showed significant sequence identity with published virus-like agents. We used a dot-blot hybridisation assay with oligonucleotide probes representing each polarity of each of ten candidate contigs and found one set which differentiated some symptomatic trees from axenically grown disease-free *E. camaldulensis*. This set of probes represented a contig of 766nts which had 95% sequence similarity to genomic DNA of *Leifsonia* bacteria (Wang et al., 2016). The sequence was identified in samples from both Australian and Syrian trees.

Our samples were harvested from predominantly one species of field grown mature eucalypt, in environmentally different geographic regions, and with different symptom severities. A majority of the samples with MY-like symptoms in the field were positive in our dot-blot assays for the 766nt contig. The axenically maintained control tree confirmed that the sequence was undetectable in the healthy *E. camaldulensis* genome. Our present evidence for an association between some MY-symptomatic eucalypts and a *Leifsonia* species suggests that a fastidious vascular bacterium (Agrios, 2005) should be investigated as a possible agent of the disease. The development of xylem-microbiome-extraction methods for woody plants (Anguita-Maeso et al., 2022) and development of media for culturing fastidious bacteria are likely difficulties needing to be overcome in progressing pathogenicity tests for bacteria in MY. Nevertheless, such biological studies should be attempted since the typical slow progression of branch dieback with the production of epicormic shoots in MY could be explained by slow basipetal migration in xylem of an agent introduced via distal infection of a branch.

Chlorotic symptoms on roadside eucalypts in Europe (E. Boa; J.W. Randles, pers. comm.) resemble those described here for Syria. Since *E. camaldulensis* is a cosmopolitan species further molecular and biological tests for *Leifsonia* species should be attempted wherever eucalypts show unexplained symptoms.

## ACKNOWLEDGMENTS

We gratefully acknowledge the late Dr. F.D. Podger and the late G. Cotton for their pioneering studies on MY; R. and J. Jeffrey for continuing observations of MY in Mundulla; Dr A. Schreiber and the Australian Centre for Plant Functional Genomics for facilitating the HTS; ICARDA for hosting DH; Dr E. Boa for roadside observations in Europe.

## Author Contributions

DH and JWR selected and processed samples, UB carried out the Illumina High Throughput Sequencing and identification of candidate contigs, DH conducted hybridisation assays, AB identified the candidate *Leifsonia* sequence. JWR wrote the draft manuscript. All authors reviewed and approved the manuscript.

## Notes

### Competing Interest Statement

The authors have declared no competing interest.

